## Supplementary Information for "The nuclear speckles protein SRRM2 is a new therapeutic target molecule on the surface of cancer cells"

### Supplementary Figures

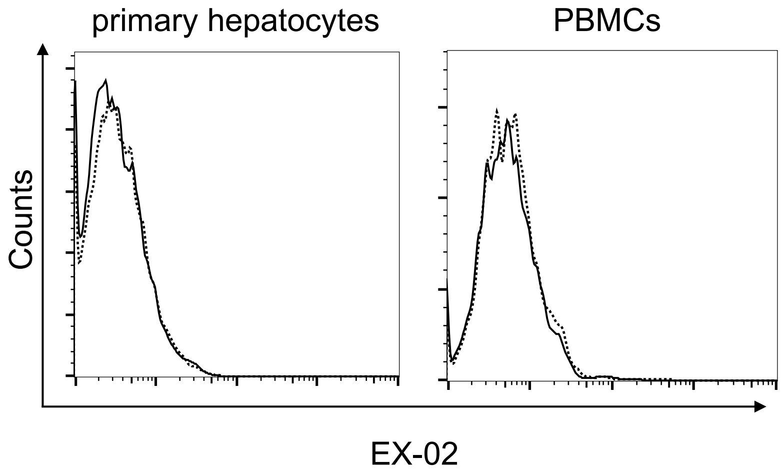


**Supplementary Figure S1:** primary hepatocytes and PBMCs stained with EX-02 and a anti-rat IgG / Alexa 647 secondary antibody. Dotted line = isotype control antibody

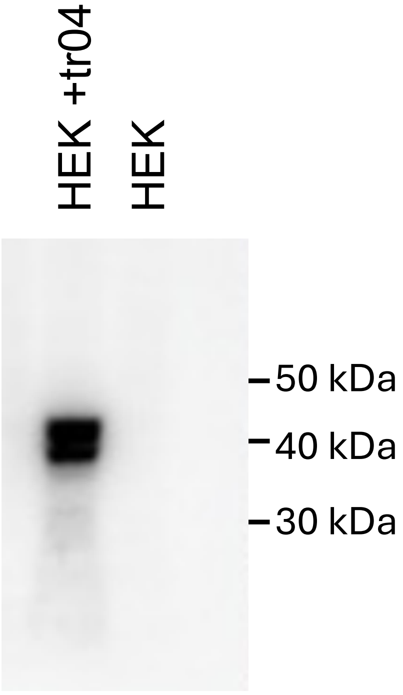


**Supplementary Figure S2:** EX-02 immunoblot. HEK293 cells were transfected with an expression plasmid encoding a tr04-his fusion protein. Lysates from transfected and non-transfected cells were separated by PAGE, blotted, and incubated with EX-02, followed by incubation with a HRP-coupled anti-rat IgG secondary antibody and finally developed with ECL.

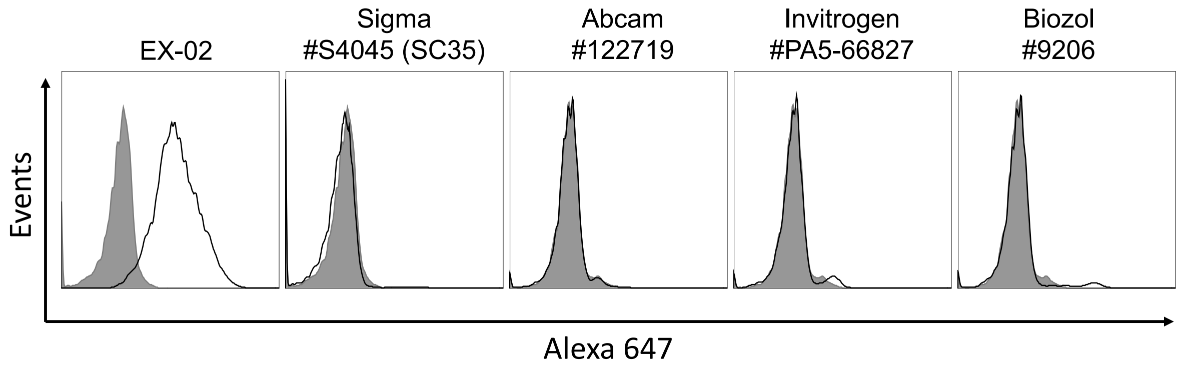


**Supplementary Figure S3A:** HeLa cells were incubated with EX-02 or commercial SRRM2 antibodies, washed, and then incubated with suitable Alexa647-labeled secondary antibodies. Binding was measured by flow cytometry. SRRM2 = black line; isotype control = tinted grey histogram.

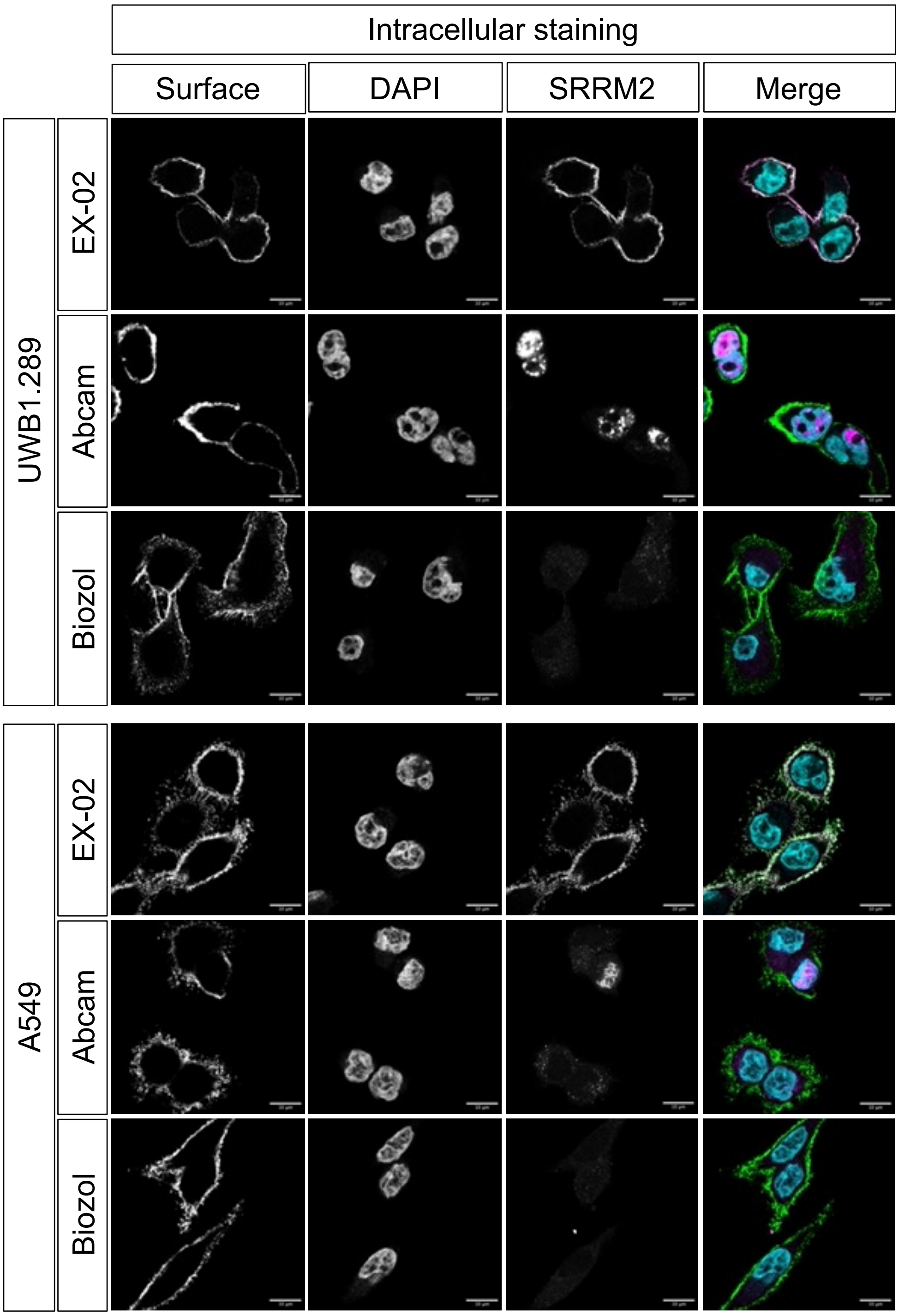


**Figure S3B:** Confocal microscopy with (A) vital and (B) fixed and permeabilized UWB1.289 and A549 cells. EX-02, Abcam, clone #122719 and Biozol, (MBS9609206) were used as SRRM2 antibodies (magenta), an EpCAM or IGF-α3 antibody were used to stain an established surface protein (green). Nuclei were counterstained with DAPI (cyan).

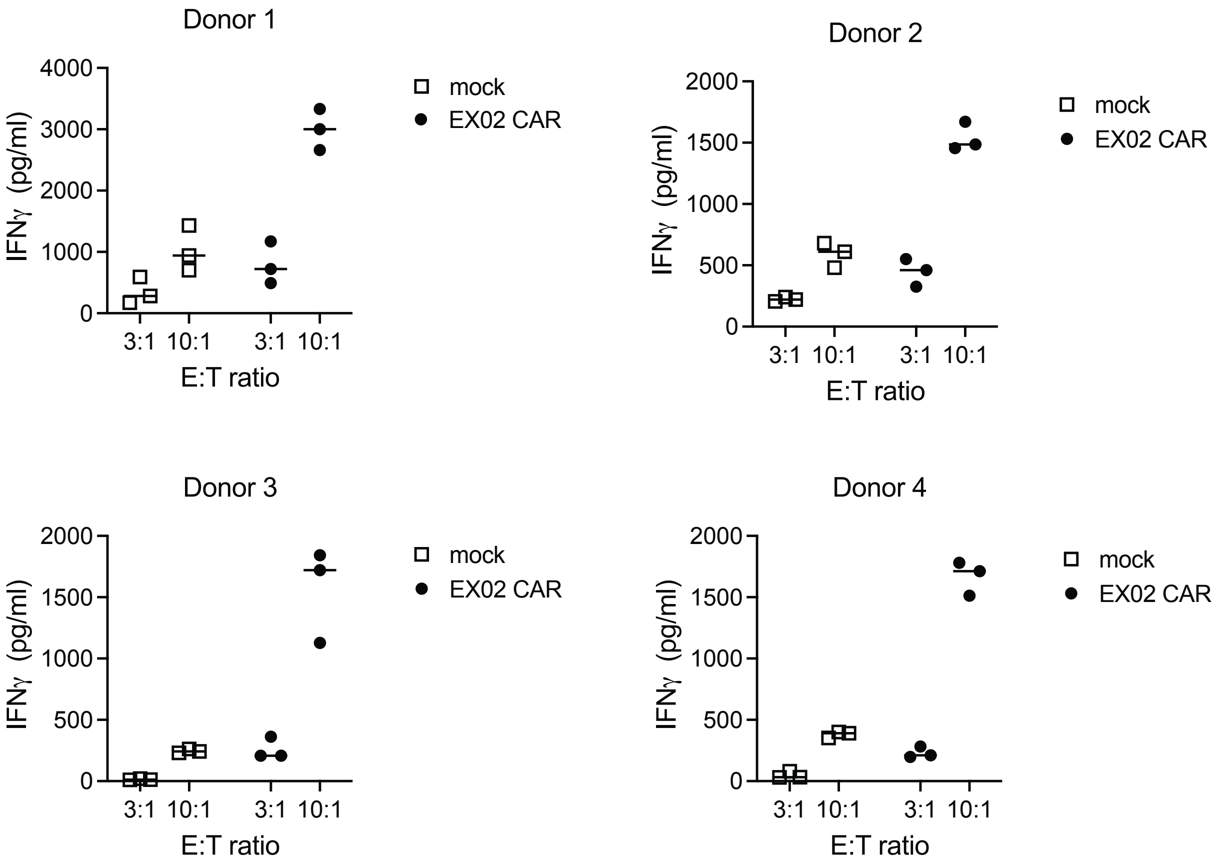


**Supplementary Figure S4:** EX02 CAR-T cells or mock-T cells obtained from four different donors were incubated with surface SRRM2 positive PCI-1 head and neck cancer cells at different ratios for 48 h. Secretion of IFN-γ into the supernatant was quantified with a commercial sandwich ELISA.

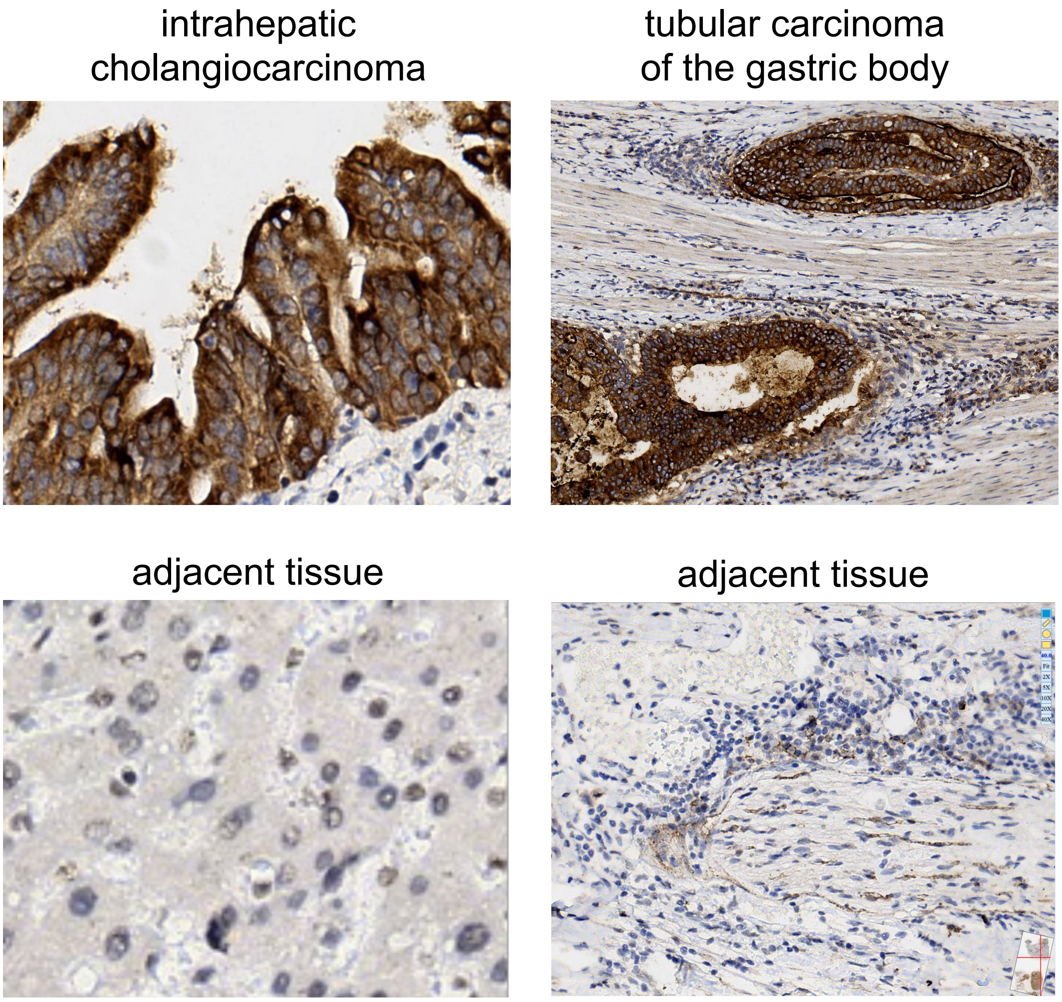


**Supplementary Figure S5:** upper panel: EX-02 membrane staining in an intrahepatic cholangiocarcinoma (left) and a tubular carcinoma of the gastric body (right), while nuclei stain almost completely negative. Lower panel: staining of adjacent non-cancerous tissues reveals weak nuclear staining.

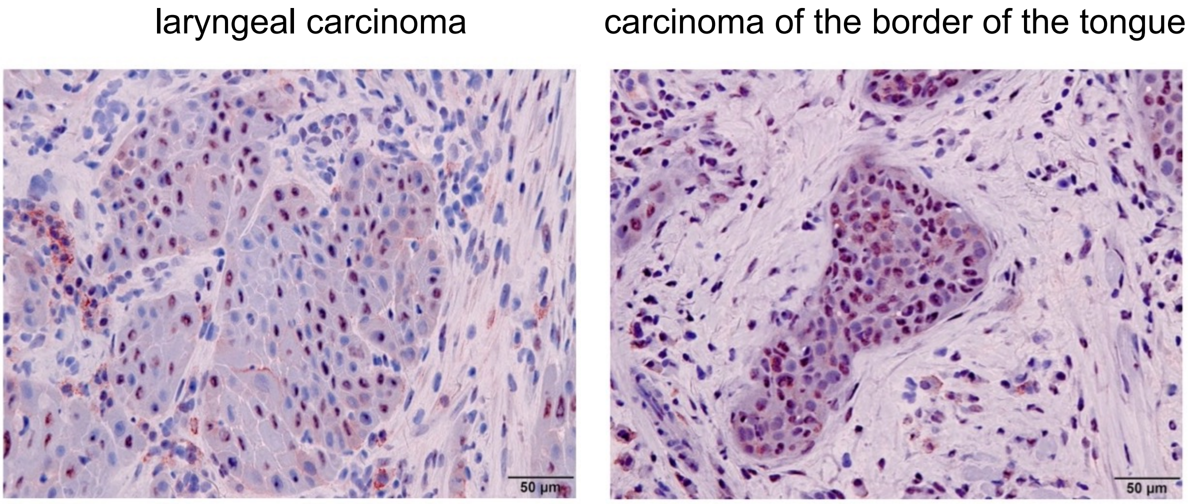


**Supplementary Figure S6:** EX-02 staining of two different squamous cell carcinoma of the larynx (left) and the border of the tongue (right) reveals nuclear localization of SRRM2 in cancer and non-cancer cells, while surface localization is not detectable.
